# Stepwise modifications of transcriptional hubs link pioneer factor activity to a burst of transcription

**DOI:** 10.1101/2022.11.08.515694

**Authors:** Chun-Yi Cho, Patrick H. O’Farrell

## Abstract

Eukaryotic transcription begins with the binding of transcription factors (TFs), which promotes the subsequent recruitment of coactivators and pre-initiation complexes. It is commonly assumed that these factors eventually co-reside in a higher-order structure, allowing distantly bound TFs to activate transcription at core promoters. Here we performed live imaging of endogenously tagged proteins, including the pioneer TF Zelda, the coactivator dBrd4, and RNA polymerase II (RNAPII), in early *Drosophila* embryos. We show that these factors are sequentially and transiently recruited to discrete clusters during activation of non-histone genes. We present evidence that Zelda acts with the acetyltransferase dCBP to nucleate dBrd4 hubs, which then trigger pre-transcriptional clustering of RNAPII; continuous transcriptional elongation then disperses clusters of dBrd4 and RNAPII. Our results suggest that activation of transcription by eukaryotic TFs involves a succession of distinct biochemical complexes that culminate in a self-limiting burst of transcription.

## INTRODUCTION

In eukaryotes, the recruitment of RNA polymerase II (RNAPII) to transcription start sites on DNA depends on the assembly of the preinitiation complex (PIC) and is regulated by hundreds of trans-acting factors^1,2^. In particular, transcription factors (TFs) recruit nucleosome remodelers, histone modifiers, and Mediator to promote the formation of PIC. How these numerous upstream inputs are integrated to give the extraordinary specificity and intricacy of transcriptional regulation remains incompletely understood. A common view suggested by biochemical studies is that these factors are progressively assembled into a single final complex through cooperative interactions. However, other sophisticated processes initiating DNA replication and promoting splicing of mRNAs are governed by a series of distinct and ephemeral complexes in which each complex promotes the next in an energy-driven step^3^. Here, we are interested in the possibility that initiation of transcription similarly involves directional transformations of intermediate complexes that would provide additional opportunity for specificity and regulation.

Visualizing the composition of transcriptional machineries over time might detect intermediate complexes that integrate the multitude of regulatory inputs of transcriptional control. In recent years, advances in confocal and super-resolution imaging led to the discovery that a wide variety of transcriptional regulators are recruited to form clusters at active genes^4^. These clusters are thought to function as “transcriptional hubs” by locally enriching transcriptional machinery and enhancing their binding to target DNA sites. Transcriptional hubs are a type of membraneless compartments, whose formation typically involves the multivalent interaction between intrinsically disordered regions (IDRs)^5^. Accordingly, IDRs are commonly found in the activation domains of TFs as well as the C-terminal domain (CTD) of Rpb1 in RNAPII^6^. Similar to the idea that a single final complex is assembled on the DNA to initiate transcription, it has been proposed that the heterotypic interactions between IDRs can give rise to a compartment that simultaneously enriches TFs, coactivators, Mediator, and RNAPII at promoters^7^. Nonetheless, how transcriptional hubs are regulated and whether they undergo compositional changes are still unclear.

Studying the dynamics of transcriptional hubs in living cells is complicated by the discontinuous and stochastic nature of eukaryotic transcription, a phenomenon also known as bursting^8^. The *Drosophila* embryo provides a powerful context to study the timing of events upstream of transcriptional initiation. The early wave of transcription in *Drosophila* embryos is coupled to the rapid nuclear division cycles such that a few hundred genes initiate a burst of transcription about 3 min after each mitosis^9–11^. The synchrony of early nuclear cycles and real-time localization of tagged transcription factors allow one to easily track events leading to the onset of transcription. A recent study demonstrates that the post-mitotic burst of transcription in early *Drosophila* embryos follows an abrupt formation of RNAPII clusters, which disperse as nascent transcript levels increase^12^. Consistent with numerous observations, a model proposes that a large excess of RNAPII recruited prior to initiation is inefficiently converted to elongating RNAPII^12–15^. What produces this pre-transcriptional RNAPII clustering and how it is coordinated with a burst of transcription are not yet fully understood. Here, we follow events during the ∼2.5 minutes between mitotic exit and the formation of RNAPII clusters and the fate of these clusters as transcription ensues.

Zelda (Zld) is one of the pioneer TFs that promote the early wave of zygotic gene expression^16–18^. Maternally supplied Zld proteins bind to thousands of enhancers and promoters, and its binding sites exhibit increased chromatin accessibility and histone acetylation^19–23^. The depletion of maternally expressed Zld significantly reduces the level of zygotic transcription, and the embryos become highly defective at the mid-blastula transition (MBT)^16^. The transactivation domain of Zld has been mapped to an intrinsically disordered region^24^. Moreover, fluorescently tagged Zld forms dynamic hubs in the nucleus^25,26^, and previous studies suggest that Zld hubs increase the local concentration of other TFs and facilitate their binding to target DNA^25,27,28^. Knockdown of Zld reduces RNAPII “speckles” in fixed embryos^29^. While these previous studies support a model in which Zld promotes the formation of transcriptional hubs to facilitate the onset of zygotic transcription, the exact mechanism has not been determined.

In this study, we combine real-time imaging and genetic perturbation to delineate a pathway that nucleates and serially transforms transcriptional hubs to trigger initiation of transcription in early *Drosophila* embryos. We show that Zld acts through transcription coactivators, including the lysine acetyltransferase dCBP and the BET protein dBrd4, to initiate RNAPII clustering at non-histone genes. Importantly, at each step in this pathway, clusters of upstream and downstream regulators colocalize only transiently, indicating dynamic changes in the composition of the visualized transcriptional hubs and potentially of chromatin-bound complexes. We propose a model in which Zld forms numerous largely unstable hubs, some of which trigger a dCBP-dependent step that recruits dBrd4 to form hubs, a subset of which then promote RNAPII clustering to stimulate transcription. Finally, we establish that transcription feeds back to destabilize dBrd4 hubs, and has a more complex effect on RNAPII hubs, initially promoting their formation while a more prolonged period of transcription promotes dispersal of RNAPII. Negative feedback of transcription on hub stability makes the hub contribution to transcription self-limiting and could lead to cycles of RNAPII accumulation and depletion, thereby contributing to the busting feature of transcription. We suggest that initiation of transcription, like initiation of replication, involves stepwise modifications in the machinery to allow robust regulation.

## RESULTS

### The pioneer transcription factor Zelda acts with the lysine acetyltransferase dCBP to initiate RNAPII clustering

We sought to understand what triggers the abrupt formation and subsequent dispersal of RNAPII clusters during a burst of transcription in early *Drosophila* embryos^12^. As a previous study showed that the depletion of Zld reduced RNAPII speckles in immunostaining^29^, we wanted to further characterize this process using real-time approaches. To block the actions of Zld in the nucleus, we sequestered endogenously GFP-tagged Zld in the cytoplasm by the JabbaTrap, which is an anti-GFP nanobody fused with the lipid-droplet protein Jabba, and then recorded RNAPII dynamics using mCherry-tagged Rpb1^12,30,31^. During a normal cell cycle in control embryos, RNAPII abruptly formed two classes of clusters about 2-3 minutes after mitosis, including the large clusters at the two histone locus bodies (HLBs) and more numerous small clusters (Fig. 1a, top). In embryos injected with JabbaTrap mRNA, inhibition of GFP-tagged Zld in cycle 12 blocked the formation of small RNAPII clusters at non-histone genes but not the large clusters at HLBs (Fig. 1a, bottom). The timing of injection of *JabbaTrap* mRNA can be adjusted, and because the accumulated JabbaTrap only sequesters its nuclear targets during mitosis when nuclear membrane breakdown exposes Zld to the cytoplasmic trap, we achieved abrupt trapping of Zld at the transition from one cycle to the next. We found that abrupt sequestration of Zld in mitosis 12 still blocked most RNAPII clustering in the following interphase in cycle 13 (Extended Data Fig. 1). Thus, Zld is required in both cycle 12 and 13 to initiate RNAPII clustering at non-histone genes, consistent with a role of Zld in accelerating transcription after mitosis^26,27^.

**Fig 1.**
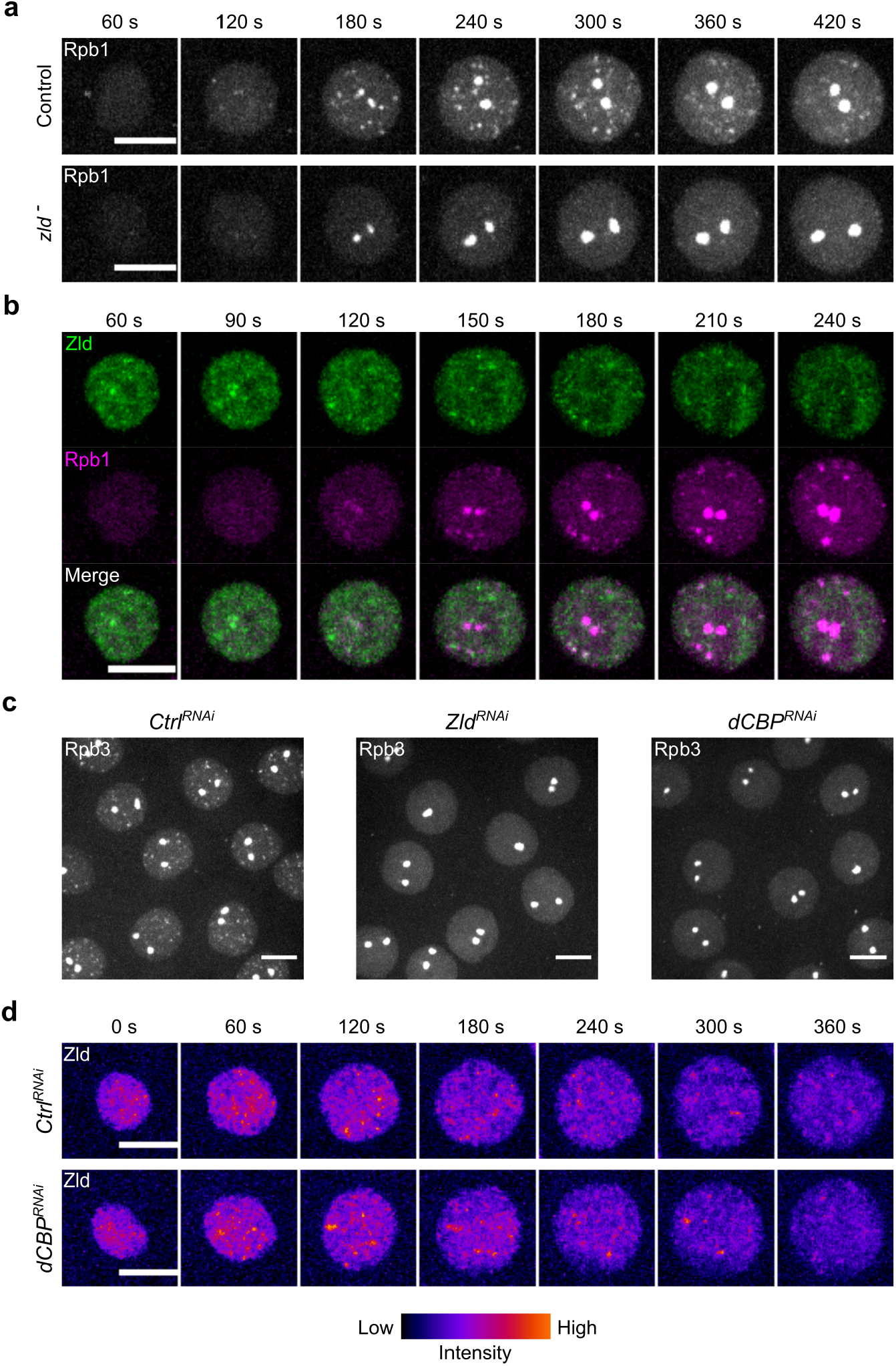
The pioneer transcription factor Zelda acts with the lysine acetyltransferase dCBP to initiate clustering of RNAPII at non-histone genes. **a**, Representative stills from live imaging of mCherry-Rpb1 in embryos with or without nuclear-localized sfGFP-Zld during cycle 12. Embryos were injected with either water (control) or JabbaTrap mRNA (*zld*^*-*^), whose protein product sequestered endogenously GFP-tagged Zld to lipid droplets (see Extended Data Fig. 1a for example). Similar outcomes were observed in 9 embryos for each treatment. **b**, Representative stills from live imaging of sfGFP-Zld and mCherry-Rpb1 during cycle 12. The timepoint 0 refers to the start of interphase. **c**, Snapshots from live imaging of EGFP-Rpb3 in embryos expressing shRNA targeting *white* (control), *zld*, or *dCBP*. Frames at 3-4 minutes after mitosis 11 are displayed. Similar outcomes were observed in at least 5 embryos for each RNAi. **d**, Representative stills from live imaging of mNeonGreen-Zld in embryos expressing shRNA targeting *mCherry* (control) or *dCBP*. Similar outcomes were observed in 5 embryos for each RNAi. All scale bars in Fig.1 indicate 5 μm.

It has been shown that Zld forms transient clusters in the interphase nucleus and that both Zld and RNAPII clusters are the most prominent before substantial accumulation of nascent transcripts^12,25,26^. We thus tested if early RNAPII clustering results from its direct recruitment to Zld hubs. We used live microscopy to simultaneously follow the dynamics of sfGFP-Zld and mCherry-Rpb1 (Fig. 1b). Upon entering interphase in cycle 12, Zld clusters emerged promptly within a minute, while general nuclear RNAPII gradually increased but was still uniformly distributed during this period (Fig. 1b, 60-120 seconds). Shortly thereafter, RNAPII clusters emerged but only infrequently overlapped with a small subset of Zld clusters (Fig. 1b, 150-180 seconds). When RNAPII clusters became more prominent (Fig. 1b, 180-240 seconds), the overall intensity of Zld clusters decreased, and the remaining faint Zld foci rarely colocalized with the matured RNAPII clusters. These observations are not consistent with the recruitment of RNAPII by direct interaction with Zld, which would predict high spatial and temporal correlation of their fluorescent intensities. Instead, these findings suggest intermediate steps between Zld and RNAPII clustering before the onset of transcription.

Previous work suggested events that might mediate an indirect requirement of Zld for RNAPII recruitment. Zld binding increases chromatin acetylation and DNA accessibility^19–23^, but the coactivators that act with Zld remain unknown. Among the chromatin changes associated with Zld, the acetylation of histone H3 lysine 27 (H3K27ac) is known to be deposited by the lysine acetyltransferase CBP/p300 and is a major determinant of enhancer activity^23,32^. We thus tested whether the *Drosophila* homolog of CBP, encoded by *Nejire* (*Nej*), is necessary for RNAPII clustering. Because dCBP/Nej is essential during oogenesis such that the germline clones of the null allele do not produce eggs^33,34^, we used RNAi to deplete maternal dCBP in early embryos by expressing shRNA targeting dCBP only from the late stages of oogenesis (Extended Data Fig. 2a)^35,36^. Immunostaining confirmed the global reduction of H3K27ac in dCBP-knockdown embryos (Extended Data Fig. 2b). Except for the large HLBs, the clustering of EGFP-tagged Rpb3 (another subunit of RNAPII) in cycle 12 was blocked by the knockdown of dCBP, phenocopying the knockdown of Zld (Fig. 1c)^21^. Knockdown of dCBP did not abolish the clustering of mNeonGreen-tagged Zld^37^ presumably at its target DNA sites, suggesting that dCBP is not required for the initial binding of Zld.

We conclude that Zld and dCBP are both required to initiate RNAPII clustering at non-histone genes and that Zld acts either upstream of dCBP or in parallel in this process. Additional evidence presented below shows that Zld and dCBP are both required for yet another intermediate step upstream of RNAPII clustering.

### Clusters of different transcriptional regulators emerge at distinct times after mitosis

We hypothesized that acetylation marks deposited by dCBP subsequently initiate RNAPII clustering via recruitment of chromatin readers, such as those recognizing acetylated lysine residues. In line with this hypothesis, the mammalian bromodomain and extraterminal (BET) protein Brd4 binds acetylated histones and is critical for the formation of transcriptional condensates at super-enhancers^38–40^. *Drosophila melanogaster* has only one BET protein similar to Brd4, encoded by *female sterile (1) homeotic*. Like its mammalian orthologs, dBrd4 functions in transcriptional regulation, as shown in cell lines and gastrulating embryos^41–44^. However, how maternally supplied dBrd4 regulates the minor wave of zygotic transcription in early embryos has not been examined, partly due to the female sterility of genetic mutants^45^.

We used CRISPR/Cas9 to tag the N-terminus of endogenous dBrd4 with either HaloTag or sfGFP. The N-terminal tag is present in both the long and short protein isoforms of dBrd4, as confirmed by western blot of sfGFP-dBrd4 from pre-MBT embryos (Extended Data Fig. 3a). Flies homozygous for HaloTag-dBrd4 or sfGFP-dBrd4 are healthy and fertile, and embryos laid by their females have a hatch rate similar to those by wild type. We then used confocal live imaging to examine the localization of tagged dBrd4 in early embryos. Using either TMR-HaloTag-dBrd4 or sfGFP-dBrd4, we observed the broad association of dBrd4 with chromosomes at various stages of the cell cycle (Extended Data Fig. 3b), similar to the behaviors of Brd4 homologs^40,46,47^. In addition to the general chromatin interaction, we observed discrete clusters of dBrd4 during most of the cell cycle except a short period at the beginning of interphase (e.g. Fig. 2a). We set out to understand the regulation and function of these clusters.

**Fig. 2.**
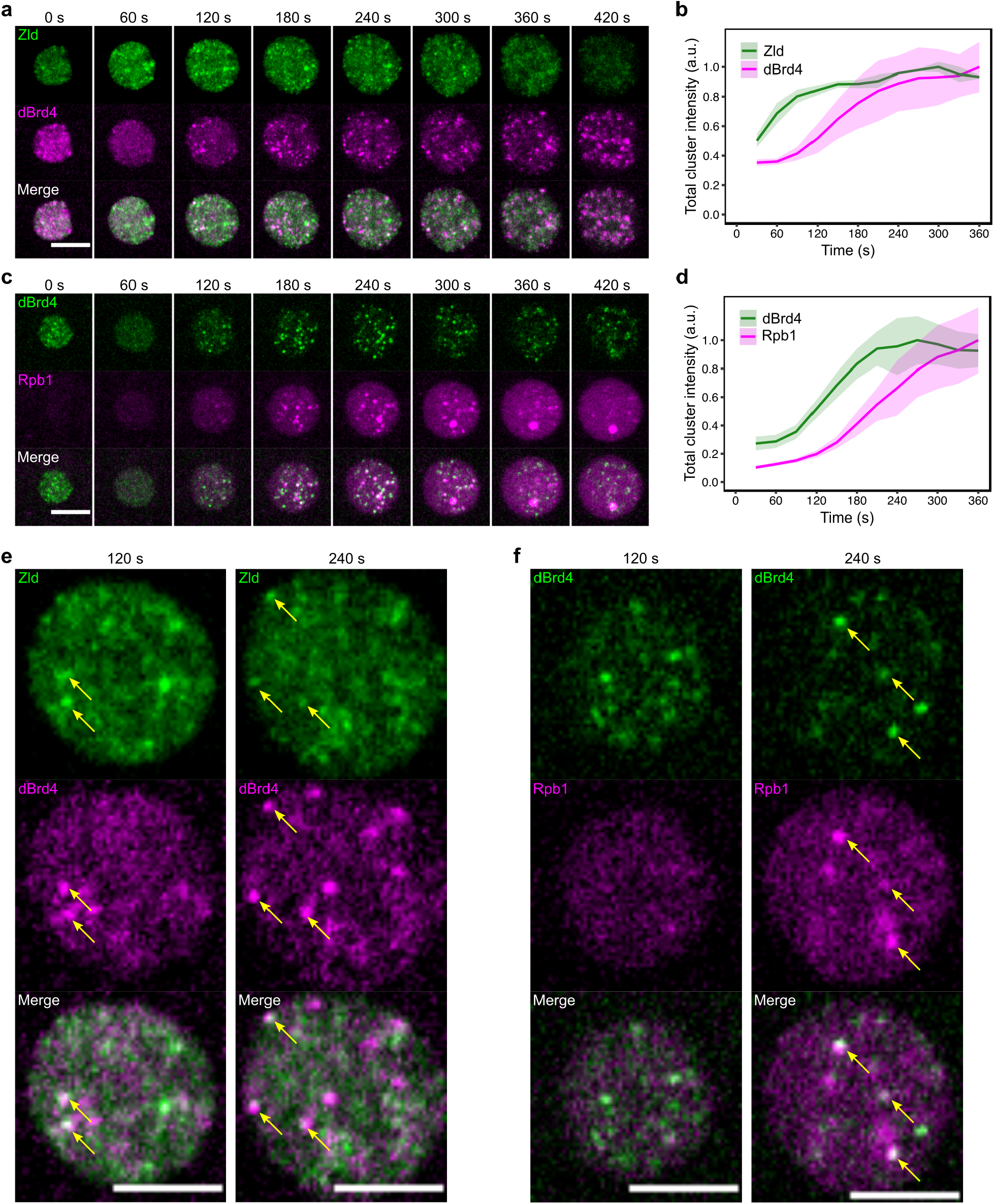
Interphase clusters of Zld, dBrd4, and RNAPII emerge sequentially after mitosis and partially colocalize with each other. **a**, Representative stills from live imaging of mNeonGreen-Zld and TMR-HaloTag-dBrd4 during cycle 12. Maximal z-projections spanning the entire nuclei are shown. Time relative to the start of interphase is indicated. Scale bar, 5 μm. **b**, Total intensities of clusters for indicated reporters per nucleus in movies as described in **a**. Clusters are defined as the pixels above 75th percentile in intensity in each nucleus. Shaded areas represent SD, n = 3 embryos. **c**, Representative stills from live imaging of sfGFP-dBrd4 and mCherry-Rpb1 during cycle 12. Scale bar, 5 μm. **d**, Total intensities of clusters for indicated reporters per nucleus in movies as described in **b**. Shaded areas represent SD, n = 3 embryos. **e**,**f**, Representative single z-plane images of indicated reporters. Time releative to the start of interphase 12 is indicated. Arrows point to examples where the clusters in two channels colocalized. Scale bars, 4 μm.

If dBrd4 clusters are involved in early zygotic transcription, their emergence in interphase might be coordinated with Zld and RNAPII recruitment. To examine this, we first performed simultaneous live imaging of mNeonGreen-Zld and TMR-HaloTag-dBrd4 in cycle 12 (Fig. 2a). Upon exiting mitosis and entering interphase, the overall chromatin-bound dBrd4 intensity rapidly decreased, and most of the mitotic dBrd4 clusters were dispersed. In contrast, Zld rapidly formed clusters from the beginning of interphase (Fig. 2a, 0-60 seconds), preceding the reestablishment of dBrd4 clusters (Fig. 2a, 120 seconds). We sought to quantify and compare the overall dynamics of clustering between Zld and dBrd4. Because both Zld and dBrd4 clusters were heterogeneous in sizes and intensities, we simply defined clusters as those pixels in the nucleus above a certain percentile in intensity, and their integrated intensity was measured after subtracting a nucleoplasmic background. Using this metric, we observed that the total intensity of Zld clusters significantly increased from 30 to 60 seconds after mitosis, whereas the total intensity of dBrd4 clusters increased more rapidly after 90 seconds (Fig. 2b). Next, we performed simultaneous live imaging of sfGFP-dBrd4 and mCherry-Rpb1 (Fig. 2c). We found that the emergence and maturation of RNAPII clusters were delayed for about 1 minute compared to dBrd4 (Fig. 2c,d). This temporal order persisted when very different intensity thresholds were used to score clusters, arguing that the measure is not sensitive to the particular parameters of the analysis (Extended Data Fig. 4). Thus, clusters of Zld, dBrd4, and RNAPII emerge in successive waves after mitosis. Moreover, visual inspection of single-z-plane images revealed that newly formed dBrd4 clusters occasionally colocalized with Zld clusters but not RNAPII, and at later times some dBrd4 clusters colocalized with RNAPII clusters (Fig. 2e,f). This suggests a model in which Zld allows dCBP to acetylate substrates and recruit dBrd4, which then mediates RNAPII clustering at Zld target genes.

### A hierarchy of genetic dependency guides the transformation of transcriptional hubs

To test whether the temporal cascade of cluster formation is also a cascade of dependency, we first used RNAi to knockdown Zld or dCBP and then examined dBrd4 localization in cycle 12. In control embryos, numerous dBrd4 clusters were visible by 3 minutes after mitosis (Fig. 3a). As predicted, following the depletion of Zld, only one to two dBrd4 clusters emerged and intensified in each nucleus after mitosis (Fig. 3a; Fig. 3b, frames 1-4). These residual dBrd4 clusters developed a signal similar to the HLBs, which normally became the dominant sites of dBrd4 recruitment in late interphase and mitosis (Fig. 3b, frames 5-8; Extended Data Fig. 5). Thus, the formation of non-HLB dBrd4 clusters depends on Zld, and it appears that an alternative pathway can promote dBrd4 recruitment to HLBs. The depletion of dCBP severely disrupted the formation of all dBrd4 clusters, suggesting that dCBP is required in both the Zld-dependent pathway for the formation of the majority of the dBrd4 clusters, as well as in the Zld-independent pathway at HLBs. Note that in addition to impairing cluster formation in interphase, the association of dBrd4 with mitotic chromosomes was reduced at non-HLB positions by Zld knockdown and was ubiquitously reduced by dCBP knockdown (Fig. 3b, frames 5-8). We conclude that dCBP is the major acetyltransferase that establishes dBrd4 clusters in early embryos. The diminishment of small dispersed dBrd4 clusters in Zld knockdown embryos suggests that Zld is one factor that can recruit dCBP, but apparently a different pathway recruits dCBP to the HLBs.

**Fig. 3.**
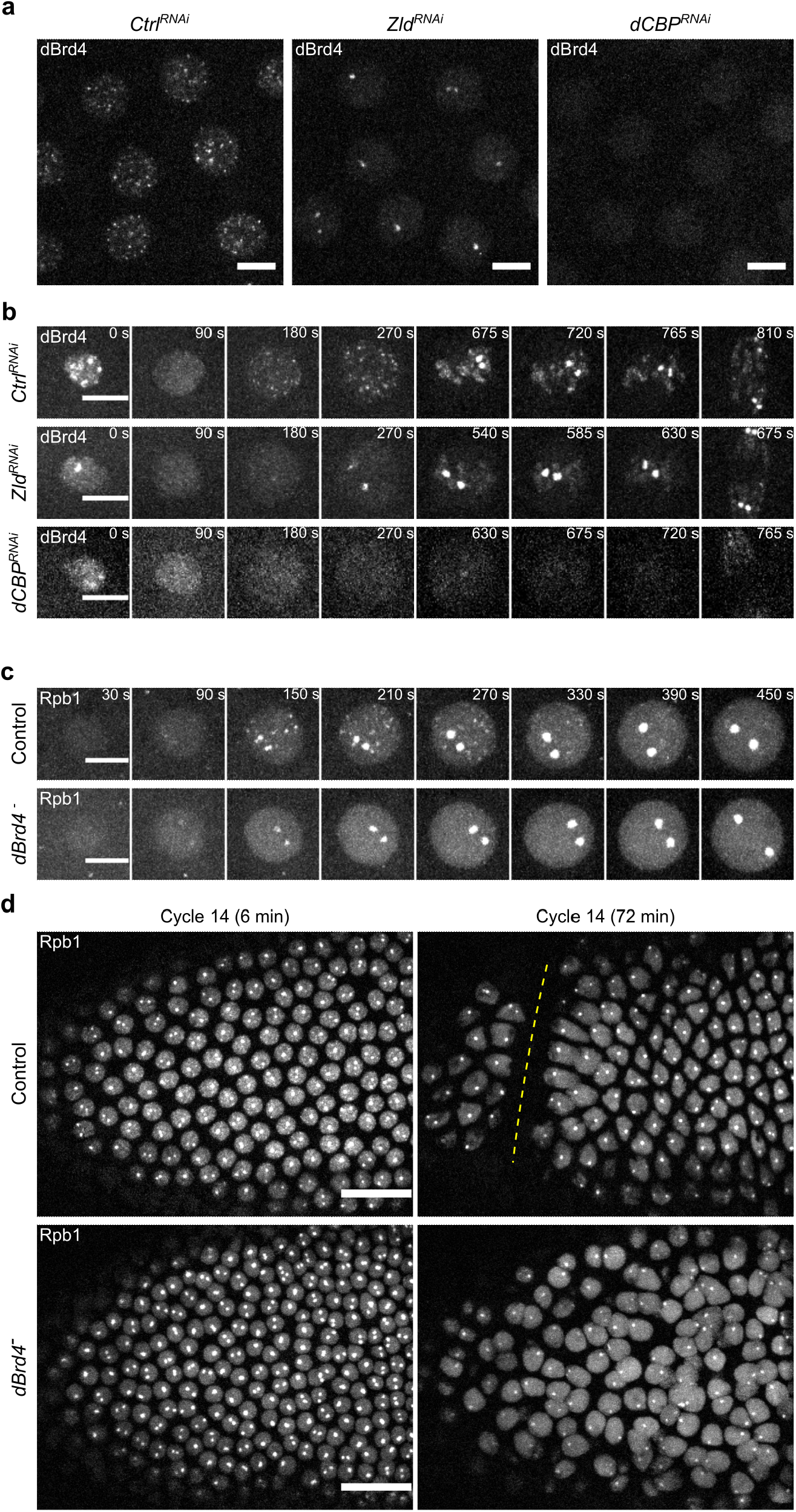
A hierarchy of genetic dependency governs the recruitment of dBrd4 and RNAPII to transcriptional hubs to activate the zygotic transcriptional program. **a**, Snapshots from live imaging of sfGFP-dBrd4 in embryos expressing shRNA targeting *white* (control), *Zld*, or *dCBP*. Frames at 3-4 minutes after mitosis 11 are displayed. Similar outcomes were observed in at least 5 embryos for each RNAi. **b**, Representative stills from movies as described in **a**. Time relative to the start of interphase 12 is indicated. The brightness and contrast for images of *dCBP* RNAi embryos were enhanced relatively to the control and *Zld* RNAi embryos. **c**, Representative stills from live imaging of mCherry-Rpb1 during cycle 12. Both control and *dBrd4*^*-*^ embryos were from females expressing *JabbaTrap* mRNA with *bicoid* 3’UTR in the germline. The untagged dBrd4 in the control embryo was unaffected by JabbaTrap, whereas endogenously GFP-tagged dBrd4 was sequestered in the cytoplasm by JabbaTrap in the *dBrd4*^*-*^ embryos. **d**, Representative stills from live imaging of mCherry-Rpb1 during cycle 14 as embryos went through cellularization and gastrulation. Anterior poles are to the left. The cephalic furrow is marked by dashed line. The genotypes for control and *dBrd4*^*-*^ are as described above. Scale bars in **a-c**, 5 μm; scale bars in **d**, 10 μm.

We next asked whether dBrd4 is responsible for RNAPII clustering. We attempted to inhibit endogenous sfGFP-dBrd4 in early embryos by maternally expressing JabbaTrap using the Gal4/UASp system; however, we found that the combination of sfGFP-dBrd4 and JabbaTrap germline expression led to female sterility, consistent with the genetic mutant phenotype^45,48^. We thus maternally expressed *JabbaTrap* mRNA with a *bicoid* 3’UTR, which includes sequences that suppress translation prior to fertilization. This allowed us to collect embryos in which sfGFP-dBrd4 was maternally expressed but was sequestered in the cytoplasm only after fertilization. In these embryos, the formation of RNAPII clusters was fully blocked except for the large clusters at HLBs (Fig. 3c). Finally, the dBrd4 JabbaTrap embryos were completely lethal and did not complete cellularization and gastrulation at the MBT (Fig. 3d), phenocopying the *zld*^*-*^ null mutant and suggesting global impairment of the early zygotic transcriptional program^16^.

We conclude that dBrd4 is the major effector of Zld and dCBP for initiating RNAPII clustering at non-histone genes.

### The spatiotemporal relationship between dBrd4 and RNAPII clusters

To gain insight into how dBrd4 triggers RNAPII clustering, we tracked individual clusters and quantified the fluorescent intensity (Fig. 4). In a typical example, we observed the earlier formation of dBrd4 clusters followed by the recruitment of RNAPII within 10-20 seconds (Fig. 4, 0-20 seconds; yellow box). The pair of dBrd4 and RNAPII clusters increased in intensity jointly for about 1 minute, and then the dBrd4 cluster began diminishing before RNAPII reached its peak intensity (Fig. 4, 60-80 seconds). The decrease in the intensity was accompanied by a disruption of cluster morphology for both dBrd4 and RNAPII. When the dBrd4 cluster disappeared, a weak and disrupted RNAPII cluster could still be observed (Fig. 4, 120 seconds). Notably, we also observed that some dBrd4 clusters emerged and disappeared without recruiting RNAPII (Fig. 4, circle), suggesting additional inputs such as non-pioneering activators or repressors.

**Fig. 4.**
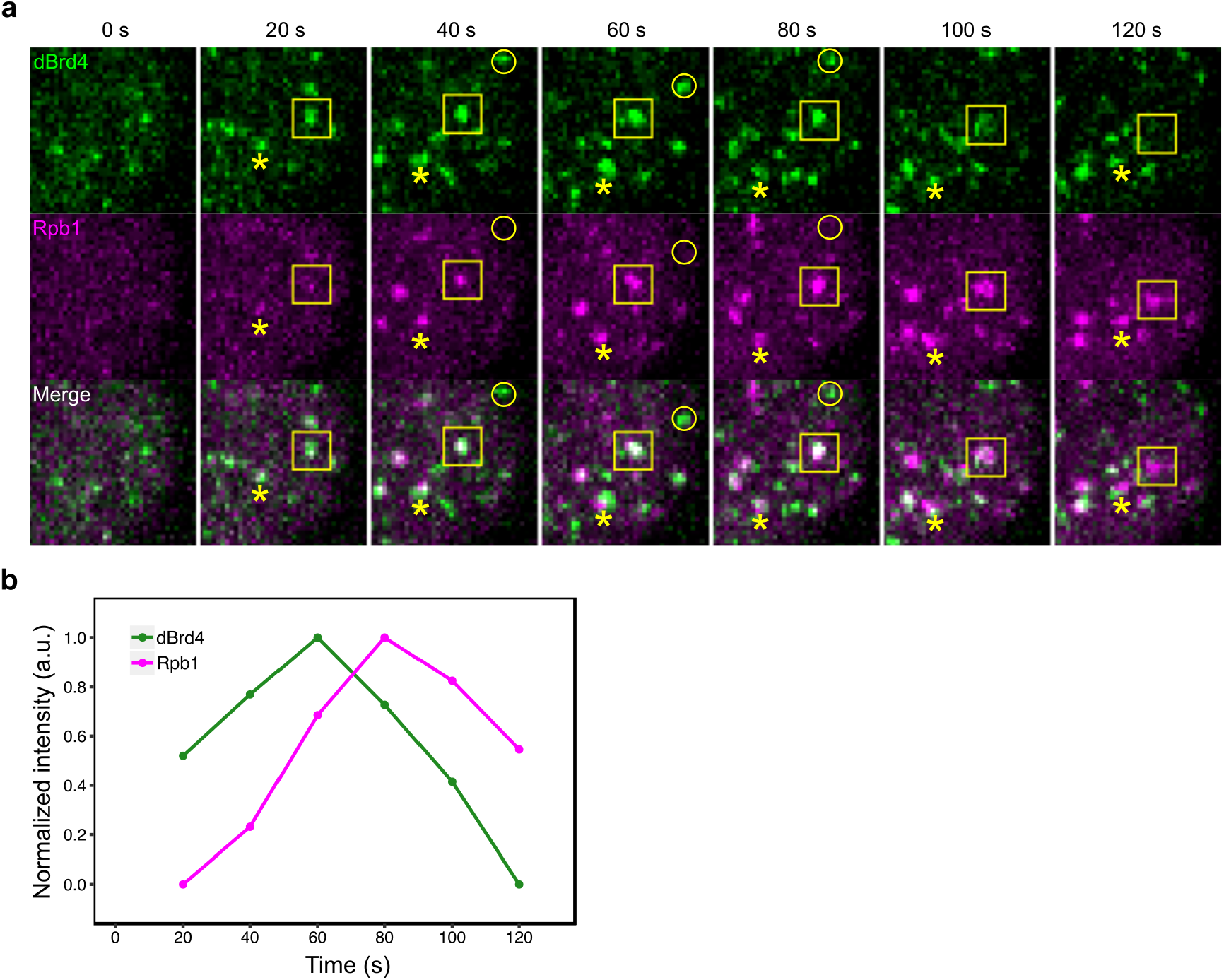
dBrd4 and RNAPII clustering are temporally separable. **a**, Stills from live imaging of sfGFP-dBrd4 and mCherry-Rpb1 during cycle 11. Maximal z-projections centered on the bottom-right portion of a nucleus are shown. Time relative to the earliest detection of dBrd4 clusters in this nucleus is indicated. The box tracks a pair of dBrd4 and RNAPII clusters as they sequentially emerge and disperse in the movie. The asterisk marks another locus that recruit both dBrd4 and RNAPII. The circle indicates a locus or perhaps different ephemeral but nearby loci where dBrd4 clustering is not followed by RNAPII recruitment. Scale bar, 2 μm. **b**, Line graph showing the normalized intensity profiles of dBrd4 and Rpb1 at the cluster outlined in the yellow box in **a**.

Unexpectedly, we frequently observed a small displacement of RNAPII clusters from associated dBrd4 signals in the live-imaging data (Fig. 4 and Extended Data Fig. 6a). Because part of this displacement could be due to movement during imaging of the two channels (less than 1 second), we sought to verify this finding in fixed embryos. We injected 2% formaldehyde into embryos at about 3 minutes in cycle 12 and incubated them for another 8 minutes. The injection of formaldehyde rapidly arrested the progression of nuclear division cycles, while the sfGFP-dBrd4 and mCherry-Rpb1 clusters were immobilized and retained their fluorescence (Extended Data Fig. 6b). In these fixed nuclei, we observed a similar separation of dBrd4 and Rpb1 signals (Fig. 5a). As a control, the same protocol applied to mCherry-Rpb1 and EGFP-Rpb3 confirmed that the two subunits of RNAPII largely overlapped at the foci (Fig. 5b).

**Fig. 5.**
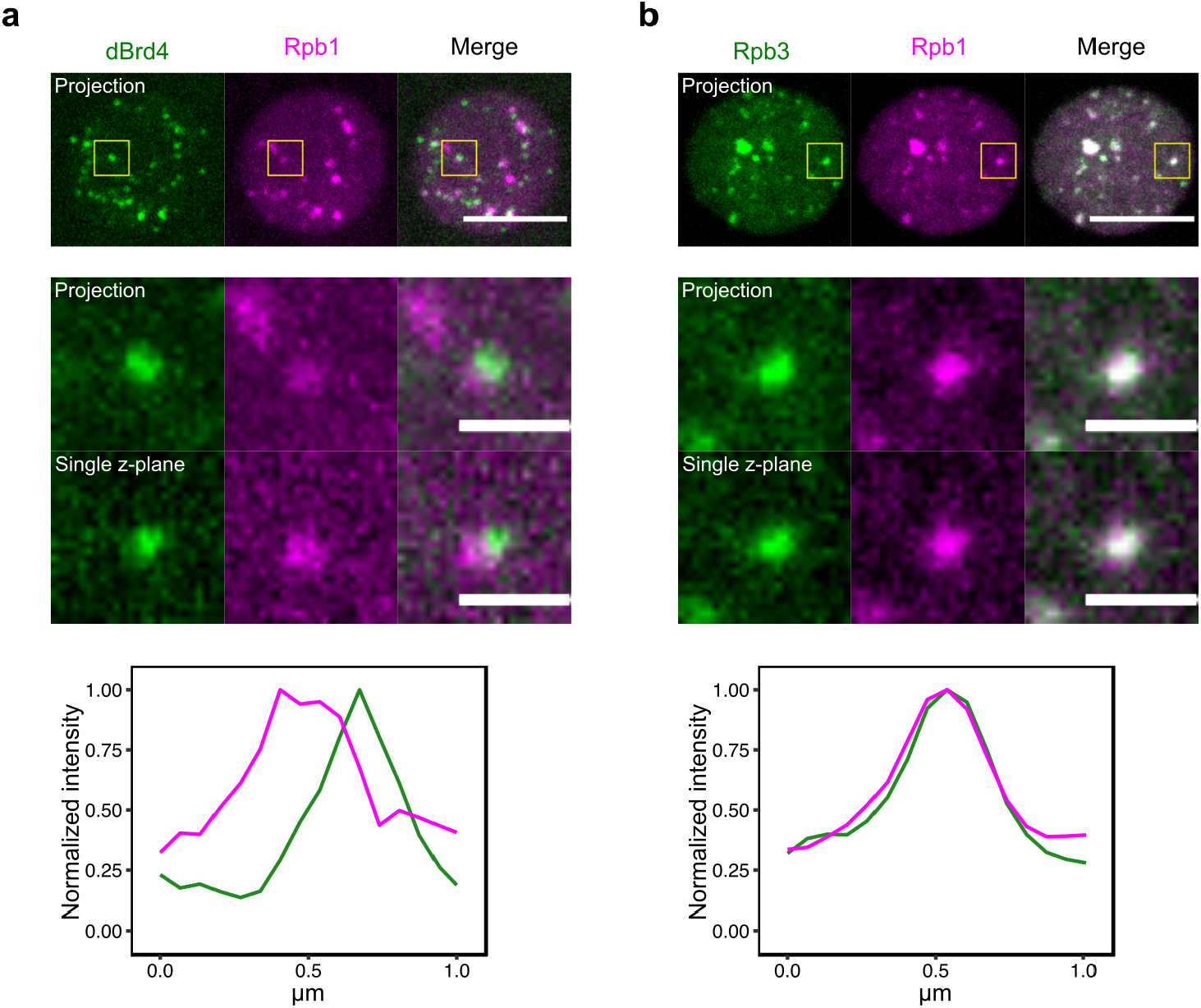
dBrd4 and RNAPII clusters are spatially separable. **a**, Snapshots from imaging of sfGFP-dBrd4 and mCherry-Rpb1 in an embryo injected with 2% formaldehyde and fixed for 8 minutes (top). The injection of formaldehyde was performed at about 3 minutes in cycle 12 to fix the clusters of dBrd4 and Rpb1. Intensity profiles from the single-z-plane image along a line perpendicular to the boundary are shown at the bottom. **b**, Snapshots from imaging of EGFP-Rpb3 and mCherry-Rpb1 in an embryo injected with 2% formaldehyde as described above. Intensity profiles from the single-plane image along a horizontal line across the cluster are shown at the bottom. Scale bars in the whole-nuclei images indicate 5 μm, and scale bars in the magnified images indicate 1 μm.

We conclude that the clusters of dBrd4 and RNAPII are separable from each other both temporally and spatially, arguing against a simple model that dBrd4 condensates partition free RNAPII. We suggest that dBrd4 clusters established at upstream regulatory regions provide a surface to nucleate RNAPII clusters.

### A sustained period of transcription disperses dBrd4 and RNAPII clusters

The above data along with previous studies support a model in which the stepwise modifications of transcriptional hubs ultimately produce locally concentrated pools of free RNAPII to stimulate transcriptional initiation. The disruption of both dBrd4 and RNAPII clusters shortly after their maturation led us to ask whether transcriptional engagement of associated genes affects the progressive transformation of transcriptional hubs.

We used a pharmacological inhibitor of RNAPII, α-amanitin, to block translocation and RNA synthesis. We injected α-amanitin before mitosis 11 and performed live microscopy in cycle 12. Inhibition of transcription by α-amanitin had no noticeable effects on the dynamics of Zld clusters in cycle 12 (Extended Data Fig. 7a). Interestingly, in the presence of α-amanitin, dBrd4 clusters still emerged normally in early interphase but then continued to grow without being dispersed, resulting in much more intense clusters in late interphase (Fig. 6a). In contrast, α-amanitin significantly delayed and reduced RNAPII clustering (Fig. 6a). These results indicate that transcription is dispensable for Zld and dBrd4 clustering but is required for the later decay of dBrd4 clusters. Since transcription is also needed for rapid recruitment of RNAPII, the stabilization of dBrd4 clusters by α-amanitin might reflect a direct contribution of transcription in destabilizing dBrd4 foci and/or a role of RNAPII in the destabilization. In any case, the effects of transcriptional inhibition indicate that transcription normally makes two contributions to the maturation of transcriptional hubs, promoting the dispersal of dBrd4 and the accumulation of RNAPII.

**Fig. 6.**
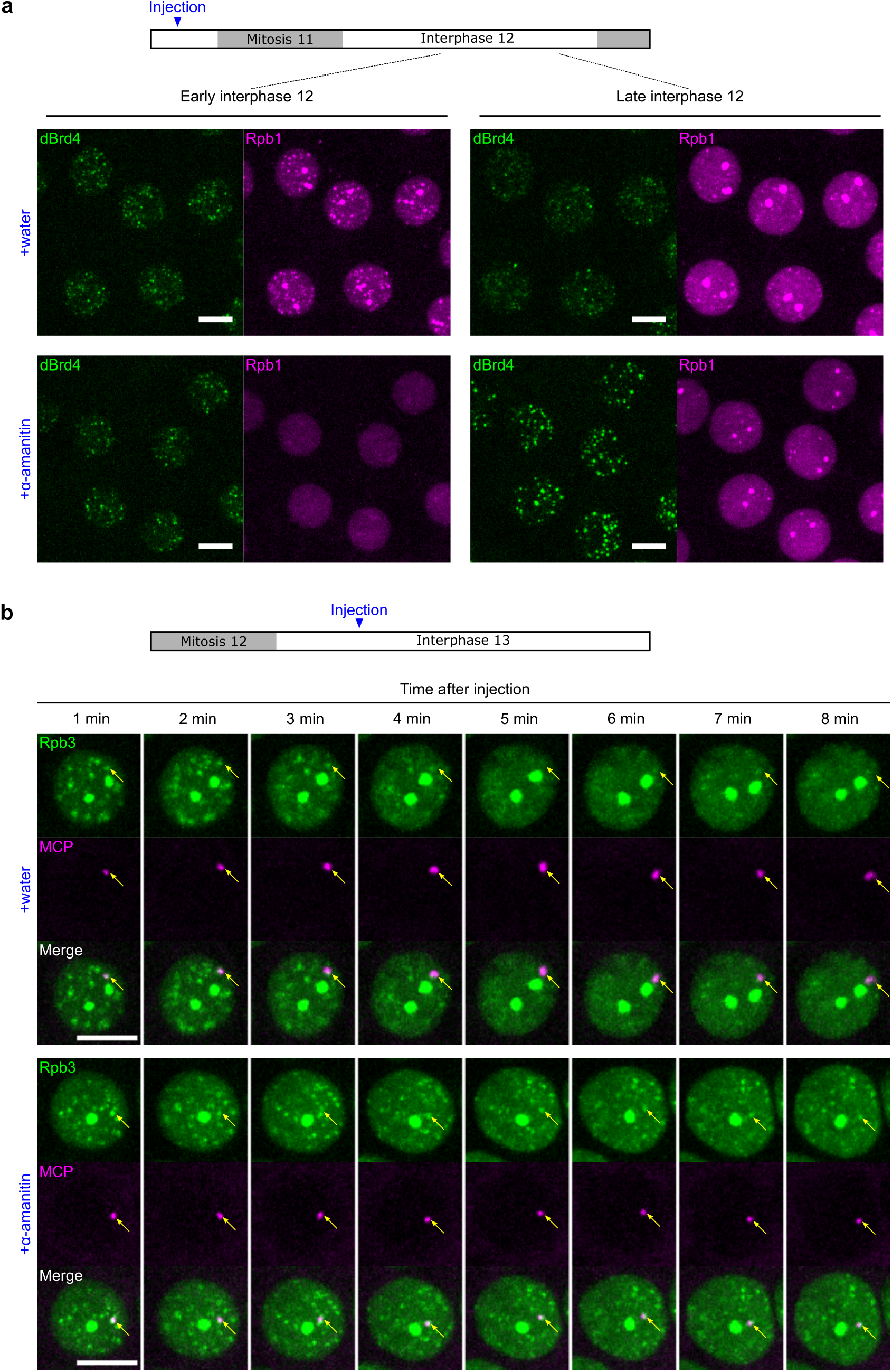
A sustained period of transcription disperses dBrd4 and RNAPII clusters. **a**, Snapshots from live imaging of sfGFP-dBrd4 and mCherry-Rpb1 at 3 minutes (early) or 6 minutes (late) in cycle 12 in embryos injected with water or α-amanitin, an RNAPII inhibitor, before mitosis 11. Similar outcomes were observed in 6 embryos for each treatment. **b**, Representative stills from live imaging of EGFP-Rpb3 and MCP-mCherry in an embryo carrying the *hbP2-MS2* reporter. Water or α-amanitin were injected around 3 minutes in cycle 13. Similar outcomes were observed in 3 embryos for each treatment. Yellow arrows indicate positions of the MCP foci. All scale bars, 5 μm.

The finding that prior α-amanitin treatment suppressed RNAPII clustering was surprising, since RNAPII accumulation was presumed to occur upstream of transcription. To examine the timing of the transcriptional input, we injected α-amanitin right after the emergence of RNAPII clusters at about 3 minutes in cycle 13. This abrupt and later attenuation of transcription still stabilized dBrd4 clusters (Extended Data Fig. 7b, top); moreover, dBrd4 was gradually redistributed into the large dBrd4 clusters at HLBs, suggesting an exchange and competition between different clusters. More importantly, the initially formed RNAPII clusters were stabilized by abrupt injection of α-amanitin (Extended Data Fig. 7b, bottom). To test whether this stabilization of RNAPII clusters occurs in association with transcribed genes, we performed similar experiments in embryos expressing EGFP-Rpb3 and MCP-mCherry with a *hbP2-MS2* reporter^10^. In control embryos injected with water, the MCP foci colocalized with RNAPII clusters transiently in the beginning of the time course, and as MCP foci grew and expanded, RNAPII clusters were dispersed (Fib. 6b). In contrast, the abrupt injection of α-amanitin arrested the growth of MCP foci, and RNAPII clusters including those associated with MCP foci persisted without dissolving. We conclude that an early period of initial transcription is sufficient to nucleate RNAPII clusters, but that sustained transcriptional activity promotes dispersal of both dBrd4 and RNAPII clusters.

## DISCUSSION

It has long been recognized that the compartmentalization of transcriptional machinery is a fundamental aspect of eukaryotic gene control. Early cytological studies revealed discrete clusters of RNAPII and nascent transcripts, which were speculated to be stable “transcription factories”^49^. Subsequent studies show that rather than genes being recruited to stable factories, numerous factors form hubs or liquid-like condensates transiently at active genes. This leaves open the questions of what governs the dynamics of transcriptional hubs/condensates and how their emergence and dispersal are linked to transcript synthesis. In this study, we used real-time approaches to dissect upstream events in transcriptional initiation, whose timing is constrained and synchronized in early *Drosophila* embryos by coupling to the rapid cell cycles. In addition to visualizing a temporal cascade, we document a parallel cascade of dependencies. The pioneer TF Zelda acts through dCBP to nucleate transcriptional hubs, which are then sequentially modified by downstream effectors including dBrd4 and RNAPII. Our findings indicate that transcriptional hubs progress directionally through a series of intermediate states with different composition, rather than simply recruiting all the factors involved in initiating transcription progressively, and that the eventual onset of transcription feeds back to disperse transcriptional hubs. We suggest that these transitions create a cycle and underly a burst of transcription, similar to the proposal that non-equilibrium dynamics of transcriptional condensates make direct contributions to bursting^50,51^.

The dynamic nature of transcriptional hubs we described herein is distinct from the well characterized transcriptional condensates at nucleoli or histone locus bodies, which are stable compartments and incorporate multiple components^5^. The dynamic process with its multiple transitions might serve to add precision and sophistication to transcriptional control. First, transitions between discrete steps could provide proofreading steps that test the stability of intermediate complexes to filter out stochastic noise and increase regulatory specificity. Second, additional regulators might promote or prevent passage through the different transitions, thereby allowing the transcriptional hubs to integrate multiple inputs to generate the intricate spatiotemporal expression of developmental genes. In line with these ideas, our data show that the transitions from Zld clusters to dBrd4 and then to RNAPII are each associated with a decline in the number of foci, suggesting that the maturation of transcription hubs is selective at successive steps. It will be important to learn how this feature contributes to the extraordinary accuracy with which the graded and combinatorial inputs generate transcriptional outputs.

The molecular mechanisms that drive the sequential transformation of transcriptional hubs remain to be fully determined. During the first step, Zld and dCBP might directly interact with each other or undergo co-condensation^52^. Alternatively, open chromatin established by Zld could facilitate binding of additional TFs that interact with dCBP^33,53,54^. In either case, it seems likely, but not yet demonstrated, that dCBP acts by increasing local acetylation to recruit the reader dBrd4. Although dBrd4 might simply bind to histone marks such as H3K27ac, the acetylation of transcriptional machinery could also be involved in recruiting dBrd4^55^. Upon crossing a concentration threshold, dBrd4 clustering might be promoted by multivalent interactions mediated by its own IDR. In the next step, our finding that dBrd4 clusters can be spatiotemporally resolved from RNAPII clusters is not consistent with a simple model that RNAPII is partitioned into dBrd4 liquid phases through the CTD. Instead, it supports a model that RNAPII clusters form at the surface of active chromatin coated by dBrd4^56,57^. Finally, we observed both positive and negative effects of transcriptional elongation on the dynamics of transcriptional hubs. The initial requirement of transcription for RNAPII clustering might involve the upstream roles of enhancer RNAs in nucleating RNAPII clusters^51^. A sustained period of transcription mediates negative feedback to disperse dBrd4 and RNAPII clusters. This could be explained by the disruption of multivalent interaction between IDRs by the negative charge of nascent RNA^51^, but other mechanisms such as histone deacetylation are also possible.

Regardless of the molecular details, we expect that similar regulatory principles are employed by evolutionarily diverse transcription factors to mediate transcriptional activation. For example, the pioneer factors Nanog, Pou5f3, and Sox19b in zebrafish similarly recruit CBP/p300 and Brd4 to establish transcriptional competence during the activation of zygotic gene expression^58,59^. Activation by estrogen receptor α (ERα) also involves histone acetylation and subsequent recruitment of BRD4^60^. Notably, elegant work has shown that dozens of factors are recruited to the ERα target promoter in a cyclical and sequential fashion^61^. We envision that many of these factors are dynamically recruited to the hubs, and that the enzymatic reactions they carry out contribute to the speed and irreversibility of the transformation of transcriptional hubs. Lastly, we suggest that the formation of transcriptional hubs in early embryos ensures the rapid initiation of a transcriptional burst within a short interphase window; in other biological contexts, the hubs might serve additional functions such as coordinating expression of multiple loci^62,63^. The *Drosophila* embryos will provide a powerful system to dissect the relationship between transcriptional hubs, chromatin interactions, and transcription dynamics.

## ACKNOWLEDGEMENTS

Stocks obtained from the Bloomington Drosophila Stock Center (NIH P40OD018537) were used in this study. We are grateful to Christine Rushlow for providing the *UASp-zld*.*shmr* fly line and to Michael Eisen for sharing the *mNeonGreen-Zld* fly line. We thank Melissa Harrison for discussing their parallel work on *nejire*. This work is supported by National Institutes of Health grant R35GM136324 (to P.H.O).

## AUTHOR CONTRIBUTIONS

**C.-Y.C**. Conceptualization, Methodology, Formal analysis, Investigation, Writing – Original Draft, Visualization. **P.O’F**: Conceptualization, Writing – Original Draft, Supervision, Funding acquisition.

---

**Extended Data Fig. 1.**
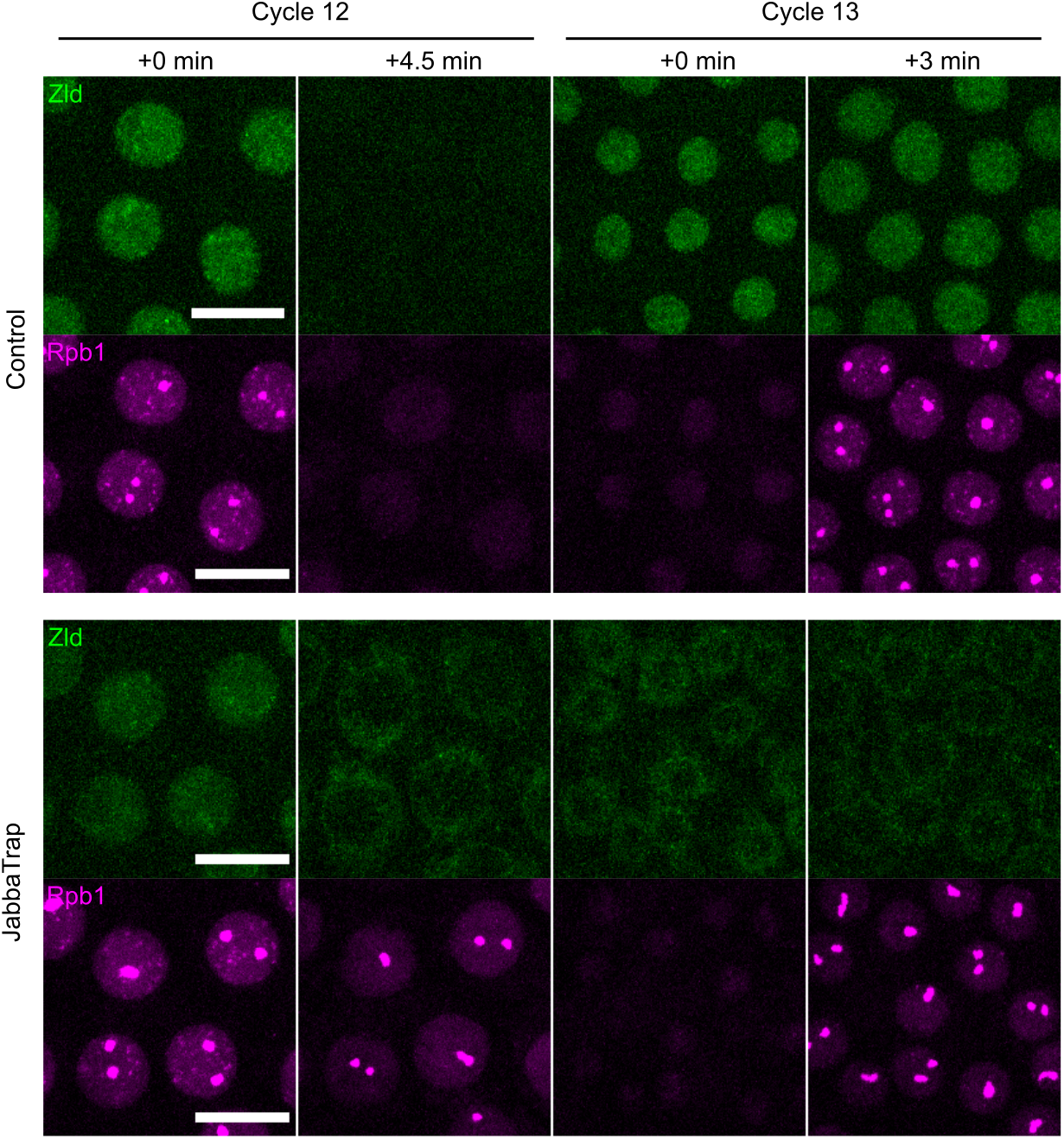
Zelda is required in cycle 13 to initiate RNAPII clustering in early *Drosophila* embryos. Live imaging of sfGFP-Zld and mCherry-Rpb1 in embryos injected either water (control) or JabbaTrap mRNA (*zld*^*-*^) to sequester endogenously GFP-tagged Zld to lipid droplets. Scale bars, 10 μm.

**Extended Data Fig. 2.**
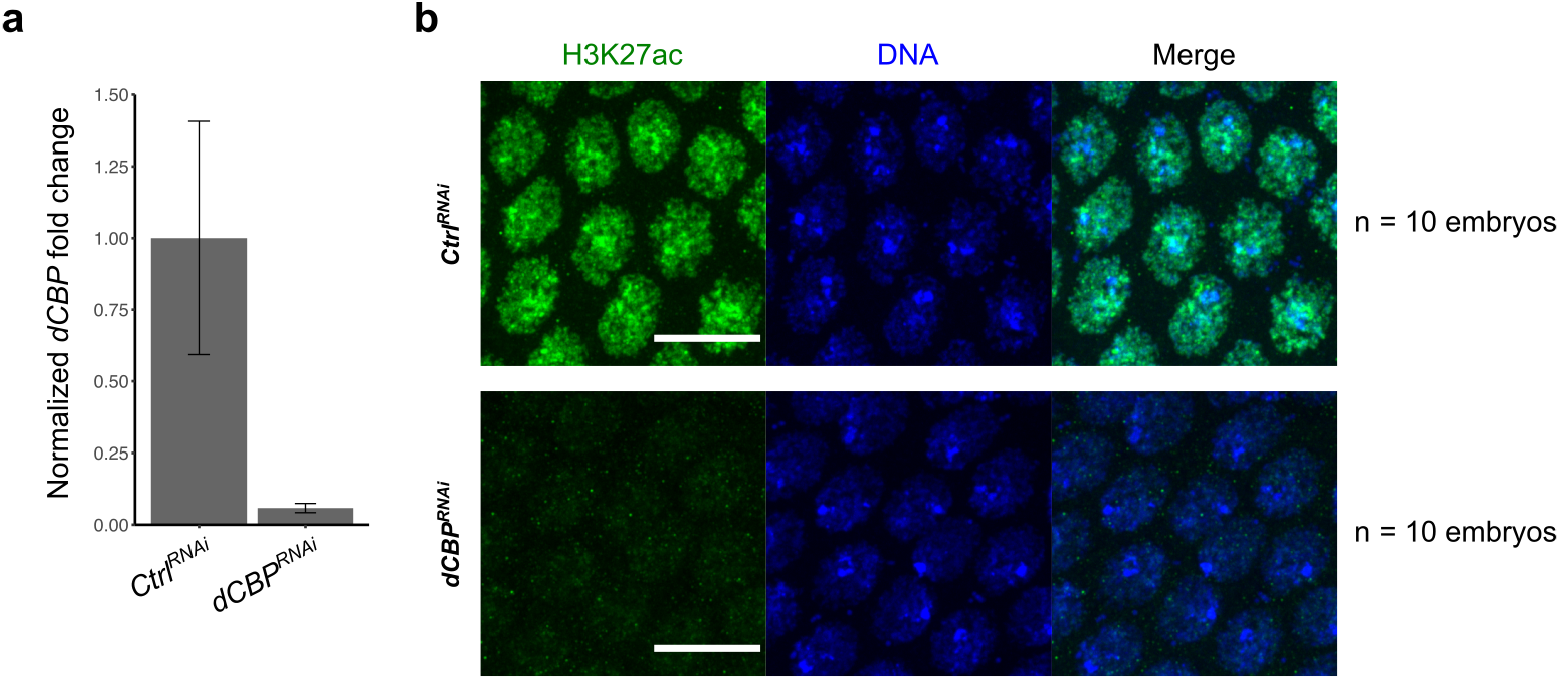
Validation of *dCBP* transcript knockdown and the reduction of histone acetylation in RNAi embryos. **a**, RT-qPCR quantification of *dCBP* transcript levels normalized by the reference gene *RpL32* in 0-1.5-hour old embryos maternally expressing shRNA targeting *mCherry* (ctrl) or *dCBP*. Error bars represent SD, n = 3 biological replicates. **b**, Representative images of H3K27ac immunostaining in embryos with RNAi knockdown for mCherry (control) or dCBP. Scale bars, 10 μm.

**Extended Data Fig. 3.**
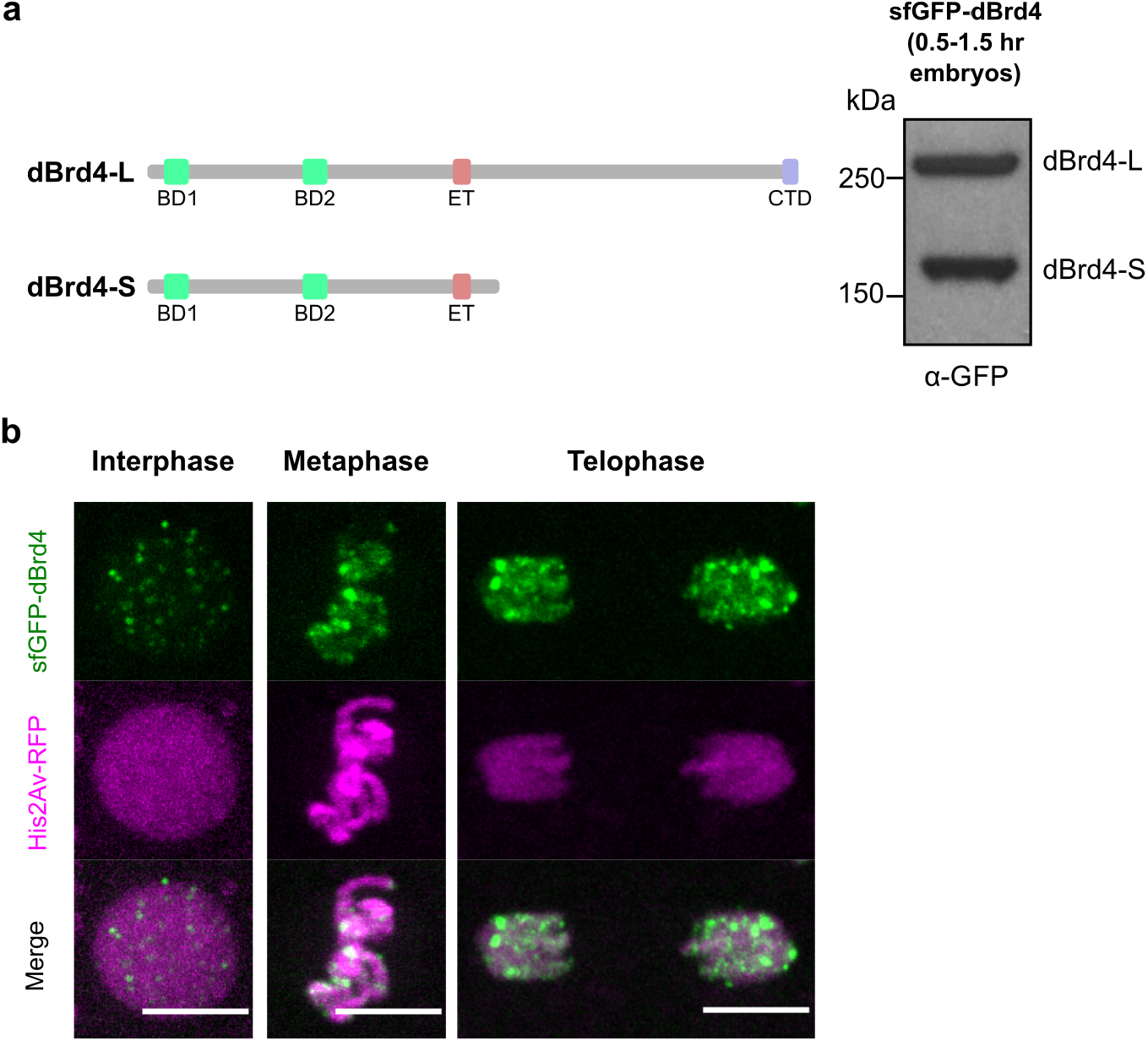
Endogenously tagged dBrd4 binds to chromatin in both interphase and mitosis. **a**, Left, schematic diagrams of the long and short isoforms of dBrd4. BD, bromodomain. ET, extraterminal domain. CTD, C-terminal domain. Right, immunostaining of sfGFP-dBrd4 from 0.5-1.5 hr embryos using an anti-GFP antibody. **b**, Snapshots from live imaging of sfGFP-dBrd4 and His2Av-RFP during indicated cell-cycle phases in cycle 12. Scale bars, 5 μm.

**Extended Data Fig. 4.**
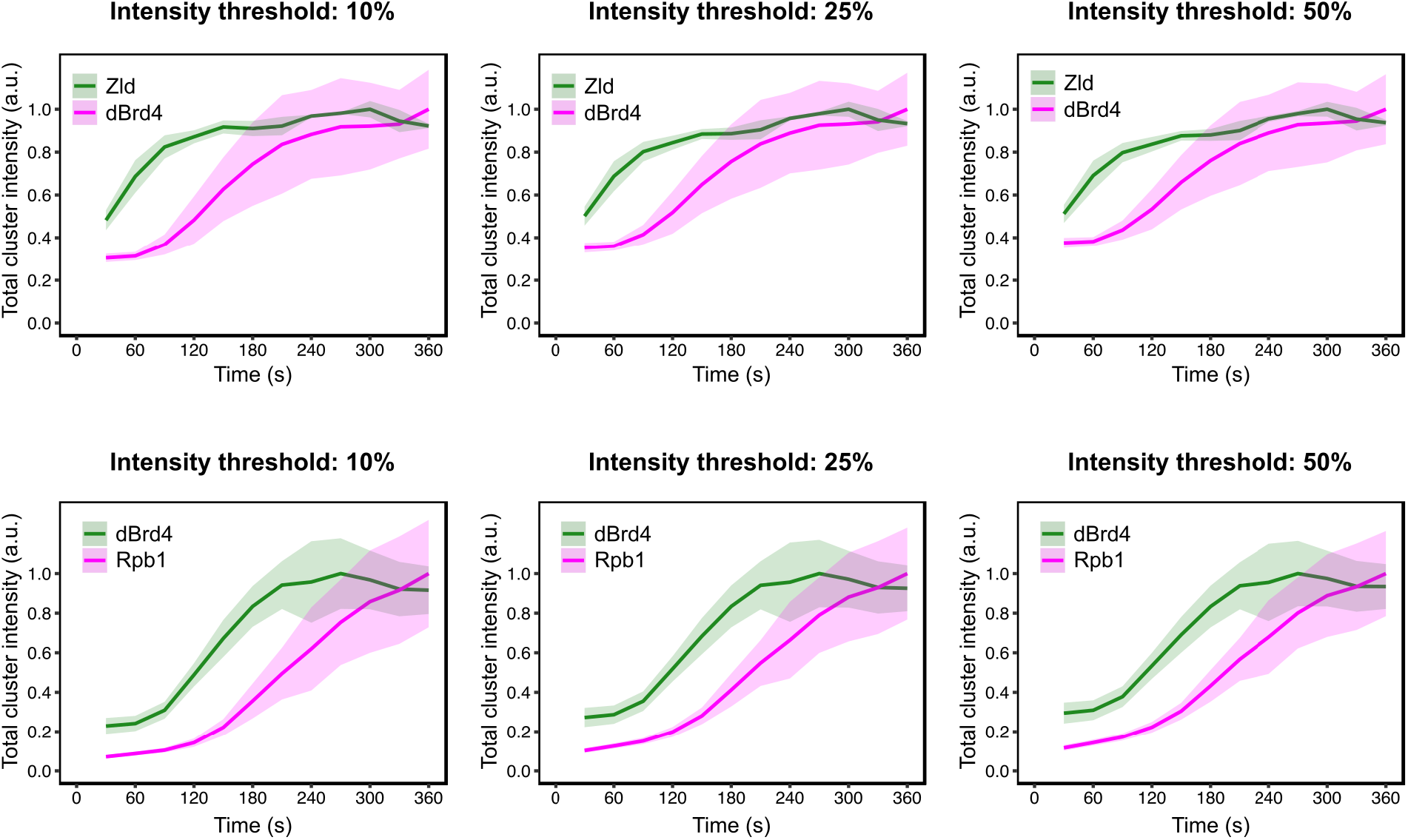
Clusters of Zld, dBrd4, and Rpb1 emerge in sequential waves during cycle 12. Line graphs showing the integrated intensities of clusters for indicated reporters during cycle 12. Clusters are defined as the pixels in the nucleus above various percentiles in intensity. Shaded areas represent SD, n = 3 embryos.

**Extended Data Fig. 5.**
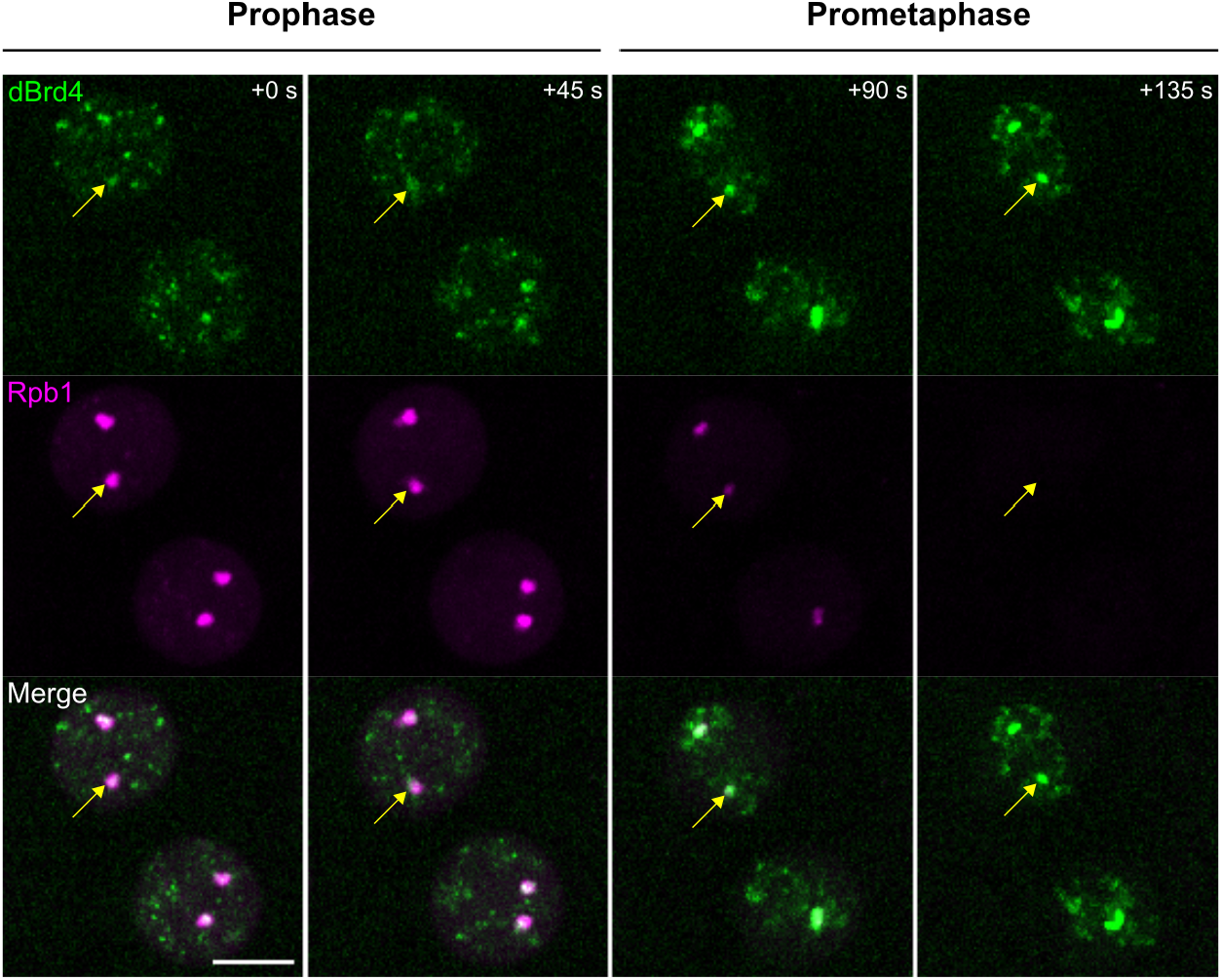
The most prominent dBrd4 clusters on mitotic chromosomes are localized at histone genes. Representative stills from live imaging of sfGFP-dBrd4 and mCherry-Rpb1 when nuclei went into mitosis 12, here visualized as the compaction of chromosomes coated by dBrd4. Time relative to the start of the movie is indicated. Arrows point to one of the histone locus bodies, initially marked by large Rpb1 clusters and later by dBrd4. Scale bar, 5 μm.

**Extended Data Fig. 6.**
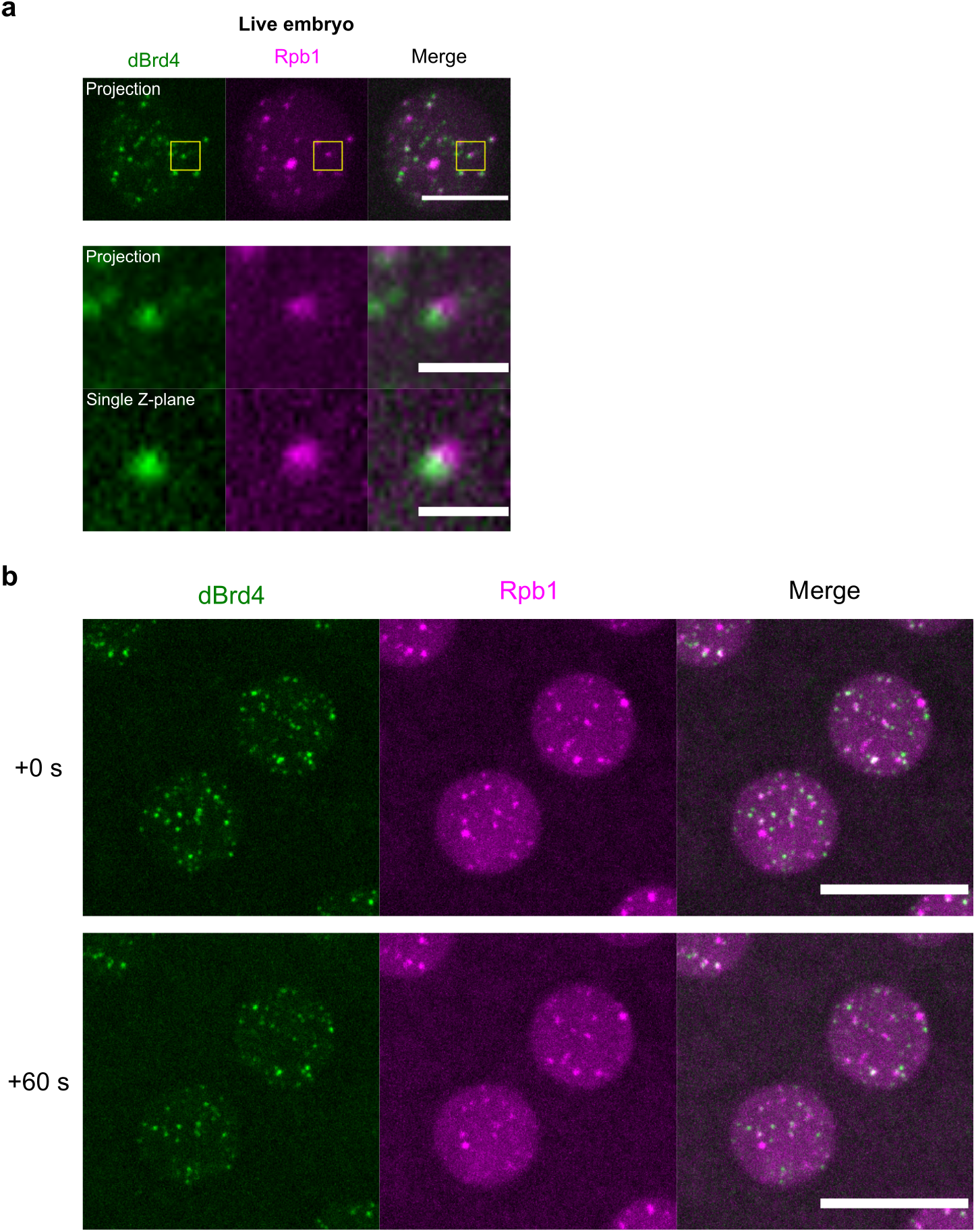
The injection of 2% formaldehyde rapidly arrests nuclear division cycle and immobilizes dBrd4 and RNAPII clusters. **a**, Snapshots from imaging of sfGFP-dBrd4 and mCherry-Rpb1 in a live embryo at about 3 minutes in cycle 12. Scale bars in the whole-nuclei images indicate 5 μm. Scale bars in the zoomed-in images indicate 1 μm. **b**, Two consecutive frames from time-lapse imaging of sfGFP-dBrd4 and mCherry-Rpb1 in an embryo fixed by formaldehyde injection. The injection was performed at 3 minutes after mitosis 11, and the embryo was further incubated for 8 minutes before taking the first frame. Maximal z-projections are shown. Scale bars, 10 μm.

**Extended Data Fig. 7.**
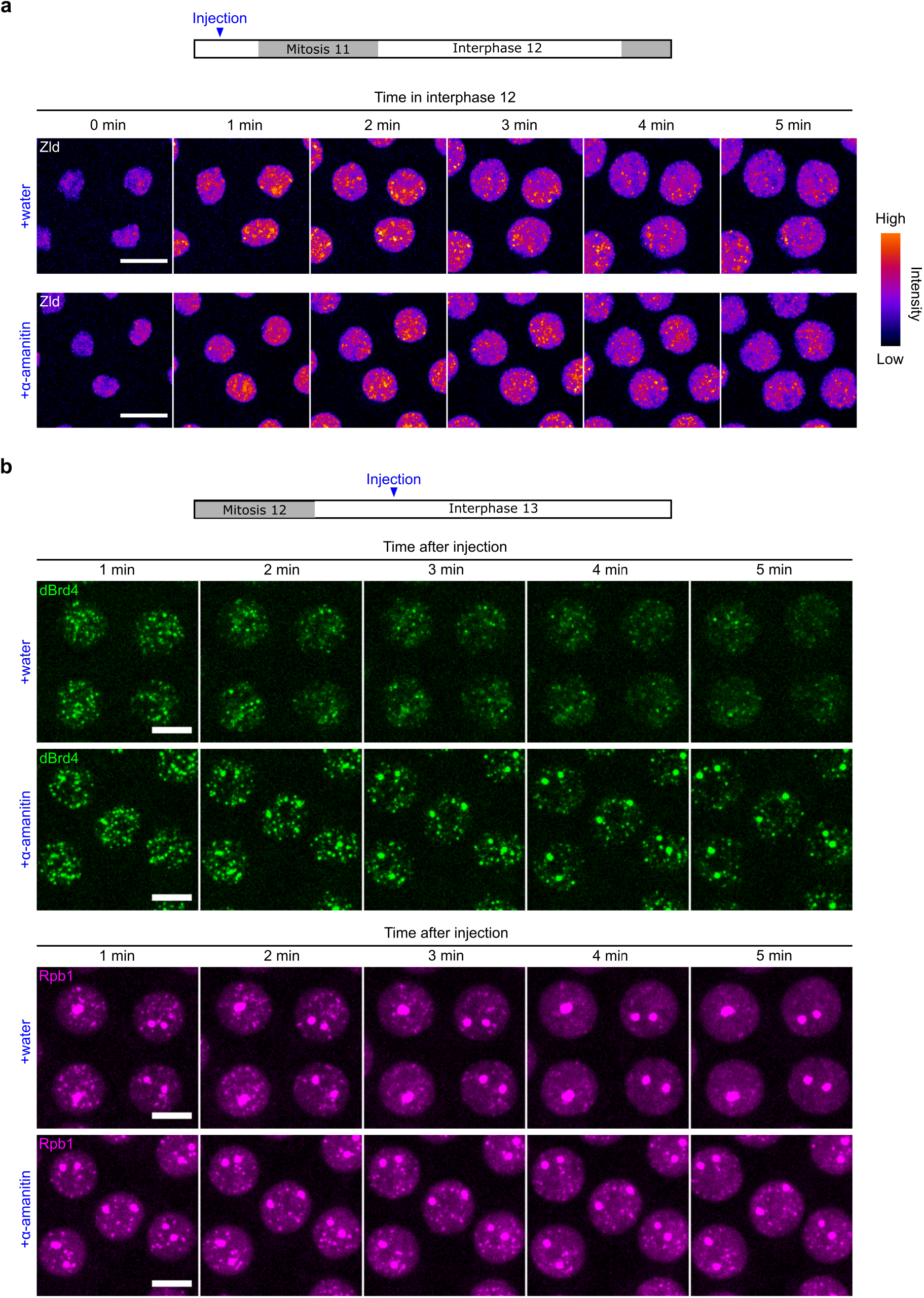
The injection of α-amanitin have different effects on clusters of Zld, dBrd4, and RNAPII. **a**, Representative tills from live imaging of mNeonGreen-Zld in embryos injected water or α-amanitin. Similar outcomes were observed in 4 embryos for each treatment. Scale bars, 8 μm. **b**, Representative stills from live imaging of sfGFP-dBrd4 and mCherry-Rpb1 in embryos injected with water or α-amanitin at about 3 minutes in cycle 13. Similar outcomes were observed in 6 embryos for each treatment. Scale bars, 5 μm.

## METHODS

### Fly stocks

*Drosophila melanogaster* were maintained on standard cornmeal-yeast medium at 25°C. Flies were transferred to egg-laying cages 2-3 days before experiments, and embryos were collected on grape juice agar plates with yeast paste. Fly lines used in this study are listed in Extended Data Table 1.

### Embryo mounting for live imaging

Embryos were collected in egg-laying cages, dechorionated with 40% bleach, washed three times with water, and then transferred onto grape juice agar plates. Embryos were aligned and transferred to coverslips with glue derived from double-sided tape using heptane. The embryos were then covered with halocarbon oil (1:1 mixture of halocarbon oil 27 and 700) for experiments.

### Microinjection

After aligned and glued to a coverslip, embryos were desiccated in a desiccation chamber for 7-9 minutes before being covered in halocarbon oil for microinjection. JabbaTrap mRNA was synthesized as described^30^ and injected at 750 ng/μl. α-amanitin (Sigma-Aldrich, #A2263) was dissolved in water and injected at 0.5-1 mg/ml. Formaldehyde was diluted fresh from 16% stock solution (Thermo Fisher, #28906) with water and injected at 2%.

### Fly crosses for germline RNAi experiments

Virgins carrying the fluorescent tag (*EGFP-Rpb3, mNeonGreen-Zld*, or *sfGFP-dBrd4*) and *UASp-shRNA* transgenes were crossed to males carrying *Mat-tub-Gal4* with or without corresponding fluorescent tag (see Extended Data Table 1 for details). The resulting progenies were then crossed with their siblings and transferred to cages for experiments. To increase the Gal4/UASp induction, F1 progeny was grown at 27°C since larval stage.

### CRISPR-Cas9 genome editing

Oligos for sgRNA targeting 5’ ends of *dBrd4*/*Fs(1)h* (5’-CGGTGGCTCACTGGACGACA-3’) were ordered from IDT, annealed and cloned into pU6-BbsI-chiRNA using standard protocol^64^. To make donor plasmids, about 1 kb of homology arms upstream and downstream of start codons were amplified from the genomic DNA of the *nos-Cas9* (III) flies. HaloTag and sfGFP DNA fragments with 5x Gly-Gly-Ser linker were amplified from plasmids previously made in the lab. Vector backbone was amplified from pDsRed-attP near XhoI and HindIII sites. All DNA fragments were then purified by gel purification and assembled by Gibson assembly. The sgRNA and donor plasmids were sent to Rainbow Transgenic Flies for microinjection. After injection, surviving adults were crossed to *yw, N/FM7c* balancer flies and screened by PCR for successful knock-in. Transformants were backcrossed with wild type (Canton S: *w*^*1118*^) at least three times before performing experiments.

### Labeling of HaloTag-dBrd4 in embryos

We noticed that the labeling of HaloTag-dBrd4 by fluorophore-conjugated ligand frequently caused mitotic defects, which could be alleviated in embryos from females heterozygous with untagged *dBrd4* allele. We thus used embryos from the *HaloTag-dBrd4/+* females for experiments. HaloTag TMR ligand was dissolved in DMSO at 5 mM as stock solution and diluted fresh to 10-15 μM with water for microinjection. Injected embryos were incubated for at least 10 minutes at room temperature before imaging.

### Molecular cloning and phiC31-mediated transgenesis

To construct the donor plasmid for *UASp-JabbaTrap-bcd3’UTR*, 834 bp of *bicoid* 3’UTR was amplified from the genomic DNA of wild-type flies and inserted into *pUASp-attB-JabbaTrap* plasmid backbone by Gibson assembly. The plasmid was then injected into *attP112 (III)* lines for phiC31-mediated integration by BestGene.

### Fly crosses for JabbaTrap experiments

To set up the JabbaTrap experiments for *dBrd4*, virgins of *sfGFP-dBrd4, mCherry-Rpb1 (X);; UASp-JabbaTrap-bcd-3’UTR (III)* were crossed to males of *sfGFP-dBrd4, mCherry-Rpb1 (X);; Mat-tub-Gal4 (III)*. The resulting F1 progenies were crossed with their siblings and transferred to egg-laying cages for experiments. The control cage was set up from similar crosses using strains in which *dBrd4* was untagged. F1 progeny was grown and kept at 25°C for experiments, as growing at 27°C still led to complete female sterility.

### Embryo fixation and immunostaining

Dechorionated embryos were transferred into a 1.5-ml tube with 500 μl heptane and then added with 500 μl of fresh 4% formaldehyde in PBS and then vigorously shaken for 20 minutes at room temperature. After removing the lower aqueous layer carefully, 500 μl methanol was added, and the mixture was shaken for 1-2 minutes. Devitellinated embryos that sank to the bottom were kept and washed with methanol for three times. Fixed embryos were stored at -20°C in methanol until use. To perform immunostaining, fixed embryos were rehydrated with 500 μl PBST (0.3% Tween-20) for 5 minutes at room temperature four times. Embryos were blocked in PBST with 3% donkey serum (Sigma-Aldrich) for 30 minutes. 1:500 rabbit α-H3K27ac (Active Motif, #39133) was then added and incubated at 4°C overnight. Next, embryos were washed with PBST for 15 minutes three times, incubated with 1:500 anti-rabbit Alexa Fluor 488 (Thermo Fisher, #A11008) at room temperature for 1 hour, washed with PBST for 15 minutes three times, and mounted on a glass slide in Fluoromount (Sigma-Aldrich, #F4680). Hoechst 33258 (Thermo Fisher, #H3569) was added at 1:2000 during the second wash after secondary antibody incubation.

### Embryo RNA extraction and RT-qPCR

For each biological replicate, about 100 dechorionated embryos were homogenized by a plastic pestle in TRI Reagent, and the total RNA was extracted using Direct-zol RNA microprep kit (Zymo Research, #R2060). cDNA was synthesized using Promega GoScript Oligo(dT) (#A2790) following manufacturer’s protocol. qPCR mixture was prepared using Bio-Rad SsoAdvanced Universal SYBR Green Supermix (#172-5270) and analyzed on Bio-Rad CFX Connect system. *RpL32* was used as a reference gene, and *dCBP* level was quantified by the ΔΔCt method.

### Western blot analysis

About 100 dechorionated sfGFP-dBrd4 embryos between 0.5-1.5 hours old were transferred to 50 μl of RIPA buffer supplemented with protease inhibitors (Pierce, #A32955). Embryos were homogenized on ice by a plastic pestle, and then 50 ul of 2X SDS-PAGE sample buffer was added. After boiling for 5 minutes, 10 ul samples were separated on 7.5% SDS-PAGE at room temperature and then transferred to PVDF membrane in cold room. The membrane was blocked with TBST (0.1% tween 20) and 5% milk, blotted with 1:5000 rabbit anti-GFP (Abcam, #ab290) at cold room overnight, followed by 1:20000 anti-rabbit HRP at room temperature for 1 hour. Membrane was incubated with SuperSignal West Pico PLUS Chemiluminescent Substrate (Thermo Fisher, #34577) and exposed to film (Thermo Fisher, #34091).

### Spinning disk confocal microscopy

Imaging was performed on an Olympus IX70 microscope equipped with PerkinElmer Ultraview Vox confocal system. Movies following the mid-blastula transition were acquired with a 60x/1.40 oil objective; all other movies were acquired using 100x/1.40 oil objective. Data were acquired using Volocity 6 software (Quorum Technologies). Pixel binning was set to 2×2 for imaging of live embryos and 1×1 for fixed samples. Focal planes with a 0.5-0.75 μm z-step were recorded at each timepoint. In dual-color imaging, sequential acquisition was performed through channels first then z-planes. Fluorophores were excited with 488 and 561 nm laser lines.

Appropriate emission filters were used in most cases. When imaging sfGFP-dBrd4 and mCherry-Rpb1 at high frame rate (10 s), the “fast sequential” mode without applying emission filters was used, and care was taken to make sure that there was negligible bleed through under the acquisition setting. Images in the same set of experiments were acquired using the same configuration, and laser power was calibrated using a laser power meter (Thorlabs) before each imaging session.

### Image processing and presentation

Data obtained in Volocity were exported as image stacks and background-subtracted with a rolling-ball radius between 50-75 pixels in FIJI/ImageJ. All experiments were repeated at least three times with similar outcomes, and representative images are shown in the figures.

### Image analysis

All image processing, segmentation, and quantification were performed in FIJI/ImageJ and Python.

#### Quantification of total cluster intensity

We used thresholding for image segmentation of mNeonGreen-Zld, TMR-HaloTag-dBrd4, and mCherry-Rpb1. In each pair of dual-color imaging, images merged from the two channels were used to generate binary masks corresponding to nuclei by Otsu’s thresholding after Gaussian blurring. For each channel, pixels in each nucleus were ordered based on their intensity values. We define the nucleoplasmic background as the median value and clusters as those pixels above a specified percentile. After subtracting the background, the integrated intensity of clusters normalized by the number of nuclei was measured. Results from three embryos are plotted as mean ± SD.

#### Tracking and quantifications of dBrd4 and Rpb1 clusters

Time-lapse dual-color imaging of sfGFP-dBrd4 and mCherr-Rpb1 was performed at a frame rate of 10 seconds. We tracked the colocalized clusters manually based on local nearest neighbors and the dynamics of cluster intensity. The integrated intensity was then measured in a small circle, and the data were normalized between 0 and 1 range.

**Extended Data Table 1.** *Drosophila melanogaster* lines used in this study.

|  |  |
| --- | --- |
| Genotype | Source |
| *nos-Cas9* | Bloomington Drosophila Stock Center (#78781) |
| *w, mCherry-Rpb1, sfGFP-Zld* | This paper |
| *y[1] sc[\*] v[1] sev[21]; P{y[+t7.7] v[+t1.8]=TRiP.GL00094}attP2* (expresses dsRNA for RNAi of *w* under UAS control) | Bloomington Drosophila Stock Center (#35573) |
| *y[1] sc[\*] v[1] sev[21]; P{y[+t7.7] v[+t1.8]=VALIUM20-mCherry}attP2* (expresses dsRNA for RNAi of *mCherry* under UAS control) | Bloomington Drosophila Stock Center (#35785) |
| *UASp-shRNA.zld* (expresses dsRNA for RNAi of *zld* under UAS control) | 21 |
| *y[1] sc[\*] v[1] sev[21]; P{y[+t7.7] v[+t1.8]=TRiP.HMS01570}attP2/TM3, Sb[1]* (expresses dsRNA for RNAi of *nej* under UAS control) | Bloomington Drosophila Stock Center (#36682) |
| *w; EGFP-Rpb3; UASp-shRNA.w* | This paper |
| *w; EGFP-Rpb3; UASp-shRNA.zld* | This paper |
| *w; EGFP-Rpb3; UASp-shRNA.nej* | This paper |
| *w; Mat-tub-Gal4; Mat-tub-Gal4* | Bloomington Drosophila Stock Center (#80361) |
| *w; EGFP-Rpb3, Mat-tub-Gal4; Mat-tub-Gal4* | This paper |
| *mNeonGreen-Zld; Mat-tub-Gal4* | This paper |
| *w, HaloTag-dBrd4* | This paper |
| *w, sfGFP-dBrd4* | This paper |
| *w, sfGFP-dBrd4; ; His2Av-mRFP* | This paper |
| *w, HaloTag-dBrd4, mNeonGreen-Zld* | This paper |
| *w, sfGFP-dBrd4, mCherry-Rpb1* | This paper |
| *w, sfGFP-dBrd4; ; UASp-shRNA.w* | This paper |
| *w, sfGFP-dBrd4; ; UASp-shRNA.zld* | This paper |
| *w, sfGFP-dBrd4; ; UASp-shRNA.nej* | This paper |
| *w, sfGFP-dBrd4; ; Mat-tub-Gal4* | This paper |
| *w; ; UASp-JabbaTrap-bcd3'UTR* | This paper |
| *w, mCherry-Rpb1; ; UASp-JabbaTrap-bcd3'UTR* | This paper |
| *w,mCherry-Rpb1; ; Mat-tub-Gal4* | This paper |
| *w, sfGFP-dBrd4, mCherry-Rpb1; ; UASp-JabbaTrap-bcd3'UTR* | This paper |
| *w, sfGFP-dBrd4, mCherry-Rpb1; ; Mat-tub-Gal4* | This paper |
| *y[1] w[\*]; P{hbP2-MS2-lacZ}JB38F* | Bloomington<br>Drosophila<br>Stock Center<br>(#60338) |

## Notes

### Competing Interest Statement

The authors have declared no competing interest.

